# Functional Gradients of the Cerebellum: A Fundamental Movement-To-Thought Principle

**DOI:** 10.1101/254326

**Authors:** Xavier Guell, Jeremy D. Schmahmann, John D.E. Gabrieli, Satrajit S. Ghosh

## Abstract

A central principle for understanding the cerebral cortex is that macroscale anatomy reflects a functional hierarchy from primary to transmodal processing. In contrast, the central axis of motor and nonmotor macroscale organization in the cerebellum remains unknown. Here we applied diffusion map embedding to resting-state data from the Human Connectome Project dataset (n=1003), and show for the first time that cerebellar functional regions follow a gradual organization which progresses from primary (motor) to transmodal (DMN, task-unfocused) regions. A secondary axis extends from task-unfocused to task-focused processing. Further, these two principal gradients reveal functional properties of the well-established cerebellar double motor representation, and its relationship with the recently described triple nonmotor representation. These interpretations are further supported by data-driven clustering and cerebello-cerebral functional connectivity analyses. Importantly, these descriptions remain observable at the individual subject level. These findings, from an exceptionally large and high-quality dataset, provide new and fundamental insights into the functional organization of the human cerebellum, unmask new testable hypotheses for future studies, and yield an unprecedented tool for the topographical, macroscale interpretation of cerebellar findings.

## INTRODUCTION

Comprehending the relationship between macroscale structure and function is fundamental to understanding the nervous system and alleviating suffering in neurological and psychiatric conditions. One central principle in the study of the cerebral cortex is that macroscale anatomy reflects a functional hierarchy from primary to transmodal processing^1,2^. For example, higher-level aspects of movement planning and decision making are situated predominantly in the anterior aspects of the frontal lobe close to the primary motor cortex, while spatial attention and spatial awareness processes predominantly engage regions of the posterior parietal lobe that are closer to the primary somatosensory cortex^3^. Similarly, Wernicke’s area is closer to the primary auditory cortex while Broca’s area is closer to the primary motor cortex.

In contrast, and despite its growing importance in basic and clinical neuroscience, the central axis of motor and nonmotor macroscale organization in the cerebellum remains unknown. The cerebellum has extensive connectivity with motor and nonmotor aspects of the extracerebellar structures. In addition to anatomy, evidence from clinical, behavioral and neuroimaging studies indicates that the human cerebellum is engaged not only in motor control but also in cognitive and affective processing^4–23^. Further, structural and functional analyses have identified cerebellar abnormalities not only in primary cerebellar injury or degeneration, but also in many psychiatric and neurological diseases that degrade cognition and affect. Examples include major depressive disorder, anxiety disorders, bipolar disorder, schizophrenia, attention deficit and hyperactivity disorder, autism spectrum disorders^24^, posttraumatic stress disorder^25^, fibromyalgia^26^, Alzheimer’s disease^27^, frontotemporal dementia^27^, vascular dementia^28^, Huntington’s disease^29^, multiple sclerosis^30^ and Parkinson’s disease^31^. Unmasking the basic hierarchical principles of cerebellar macroscale organization can therefore have large impact in basic and clinical neuroscience.

The study of connectivity gradients in resting state fMRI data - an aspect of cerebellar functional neuroanatomy that remains largely unexplored - can provide critical information necessary to address this knowledge gap. The absence of intra-cerebellar anatomical connections makes it difficult to analyze intra-cerebellar progressive hierarchical relationships using anatomical techniques. Resting-state functional connectivity from fMRI data becomes, in this case, an optimal approach to interrogate functional relationships between nearby cerebellar structures which are not directly connected. Contrasting with the common practice of partitioning neural structures into discrete areas with sharp boundaries^32,33^, Margulies and colleagues^34^ provided a simple and powerful description of the “principal gradient” of resting-state functional connectivity in the cerebral cortex using diffusion map embedding. This gradient extended from primary/unimodal cortices to regions corresponding to the default mode network (DMN), confirming the primary-unimodal-transmodal hierarchical principle of the cerebral cortex^1,2^. Similarly, Sepulcre and colleagues^35^ revealed transitions from primary sensory cortices to higher-order brain systems using stepwise functional connectivity. The present study is the first to use these analyses in the cerebellum.

Here we set out to describe the principal gradients of intra-cerebellar connectivity by using resting-state diffusion map embedding. We aim to unmask the central axis of motor and nonmotor macroscale organization of the cerebellum, analogous to the fundamental primary-unimodal-transmodal hierarchical principle of cerebral cortex^1,2^. To further characterize the functional significance and implications of these continuous gradients, we aimed to analyze their relationship with discrete cerebellar parcellations including task activity maps, resting state maps, and distinct areas of motor (first=IV/V/VI, second=VIII) and nonmotor representation (first=VI/Crus I, second=Crus II/VIIB, third=IX/X)^36,37^. We took advantage of the newly available and unparalleled power of the Human Connectome Project (HCP) dataset, where each participant (n=1003) provided one full hour of resting-state data. We incorporated task activity maps (motor, working memory, emotion, social and language processing) from a previously analyzed subset of the same group of subjects^37^ (n=787). Maps of cerebellar representation of cerebral cortical resting-state networks were obtained from the study of Buckner et al.^36^, calculated in a different group of subjects (n=1000). Data-driven clustering and stability analyses were used to compare our findings with previous discrete cerebellar parcellations, as well as to validate our hypothesis-driven divisions. A supplementary analysis of cerebello-cerebral connectivity was used to validate our interpretation of asymmetries between the two motor and three nonmotor regions of cerebellar representation. Analysis of a single participant from our cohort tested the robustness of our findings at the individual level.

## 2. METHODS

All code used in this study is openly available at https://github.com/gablab/cerebellum_gradients

### 2.1 Human Connectome Project data

fMRI data were provided by the Human Connectome Project (HCP), WU-Minn Consortium^38^. We analyzed data from 1003 participants who completed all resting-state sessions (four 15-minutes scans per subject), included in the group average preprocessed dense connectome S1200 HCP release. EPI data acquired by the WU-Minn HCP used multi-band pulse sequences^39–42^. HCP structural scans are defaced using the algorithm by Milchenko and Marcus^43^. HCP MRI data pre-processing pipelines are primarily built using tools from FSL and FreeSurfer^44–46^. HCP structural pre-processing includes cortical myelin maps generated by the methods introduced by Glasser and Van Essen^47^. HCP task-fMRI analyses uses FMRIB’s Expert Analysis Tool^45,48^. All group fMRI data used in the present study included 2mm spatial smoothing and areal-feature aligned data alignment (“MSMAll”)^49^. We did not conduct any further preprocessing beyond what was already implemented by the HCP. Results were visualized in volumetric space as provided by HCP as well as on a cerebellar flat map using the SUIT toolbox for SPM^50–52^.

### 2.2 Diffusion map embedding

Diffusion map embedding methodology was introduced by Coifman and colleagues^53^, and its application to the HCP resting-state data as performed in this study is thoroughly described in Margulies et al., 2016^34^. Instead of analyzing data corresponding to the cerebral cortex^34^, the present study included only voxels corresponding to the cerebellum. We used data from the S1200 release (n=1003) instead of the S900 release (n=820). In brief, cerebellar data in the preprocessed HCP “dense connectome” includes correlation values of each cerebellar voxel with the rest of cerebellar voxels. In this way, each cerebellar voxel has a spatial distribution of cerebellar correlations (a “connectivity pattern”). Diffusion map embedding is a nonlinear dimensionality reduction technique and can be used to analyze similarities between functional connectivity based networks. As in Principal Component Analysis (PCA), diffusion map embedding results in a first component (or “principal gradient”) that accounts for as much of the variability in the data as possible. Each following component (each following gradient) accounts for the highest variability possible under the constraint that all gradients are orthogonal to each other. The final result of a dense connectome matrix PCA analysis would take the form of a mosaic; if this method was applied, each cerebellar voxel would be assigned to a particular network with discrete borders. In contrast, diffusion embedding extracts overlapping “gradients” of connectivity patterns from the initial matrix. For example, in Margulies et al., 2016^34^, gradient 1 extended from primary cortices to DMN areas, gradient 2 extended from motor and auditory cortices to the visual cortex, etc. Each voxel is then assigned a position within each gradient. In Margulies et al., 2016^34^, a voxel corresponding to a DMN area would be assigned an extreme position in gradient 1 (e.g. a value of 6.7 in a unitless scale from −5.4 to 6.9) and a middle position in gradient 2 (e.g. a value of 1.8 in a unitless scale from −3.0 to 5.7). In this way, the result of diffusion embedding is not one single mosaic of discrete networks, but multiple, continuous maps (gradients). Each gradient reflects a given progression of connectivity patterns (e.g. from DMN to sensorimotor, from motor/auditory cortex to visual cortex, etc.), each gradient accounts for a given percentage of variability in the data, and each voxel has a position within each gradient.

It is important to highlight that our initial dense connectome matrix includes the profile of connectivity of each cerebellar voxel with the rest of the cerebellum only, rather than with the rest of the brain. In this way, our analysis reflects the intrinsic organization of the cerebellum without invoking its connectivity profiles with the cerebral hemispheres or other brain structures. This approach allows the possibility of identifying cerebellar properties that might otherwise be obscured in whole-brain connectivity analyses. The latter approach would emphasize the relationship between cerebellar structures and cerebral resting-state networks, and potentially miss relevant gradients of connectivity patterns within cerebellar resting-state data.

Diffusion map embedding and task processing analyses were also performed using a 15 minutes resting-state run from a single subject. To avoid selection bias, we chose to analyze the HCP participant corresponding to the “single subject” download package of the HCP database. Resting-state smoothing of single-subject data was performed on the resulting gradients after diffusion map embedding calculations to avoid introducing artefactual correlations.

### 2.3 Task activity and resting-state network maps

Cerebellar task activity data from a subset of 787 HCP participants were analyzed in a previous study by our group^37^. Guell and colleagues^37^ provided Cohen’s d task activity maps thresholded at 0.5 (medium effect size). A sample size of 787 subjects ensures that a Cohen’s d value higher than 0.5 will be statistically significant even after correction for multiple comparisons in the cerebellum. These tasks include the following contrasts: *Movement* (tap left fingers, or tap right fingers, or squeeze right toes, or squeeze left toes, or move tongue) minus *Average* (average of the other four movements), assessing motor function^36^; *Two back* (subject responds if current stimulus matches the item two back) minus *Zero back* (subject responds if current stimulus matches target cue presented at start of block), assessing working memory; *Story* (listen to stories) minus *Math* (answer arithmetic questions), assessing language processing^54^; *TOM* (view socially interacting geometric objects) minus *Random* (view randomly moving geometric objects), assessing social cognition^55,56^; and *Faces* (decide which of two angry/fearful faces on the bottom of the screen match the face at the top of the screen) minus *Shapes* (same task performed with shapes instead of faces), assessing emotion processing^57^.

Cerebellar resting-state network maps were obtained from Buckner et al., 2011^36^. Buckner’s study applied a winner-takes-all algorithm to determine the strongest functional correlation of each cerebellar voxel to one of the 7 or 17 cerebral cortical resting-state networks defined by Yeo and colleagues^33^. This analysis used data from 1000 subjects.

### 2.4 Clustering analyses (Supplementary methods)

Clustering analyses on the resulting diffusion map embedding gradients included k-means clustering, spectral clustering, and silhouette coefficient analysis^58^ using the scikit-learn toolbox^59^. K-means separates samples in a previously specified number of clusters, minimizes the sum of the squared differences of each data point from the mean within each cluster, but makes the assumption that clusters are convex. Spectral clustering does not have a convexity constraint, provides a valuable alternative method of analysis to validate k-means clustering results, but still requires a specification of a number of clusters. Silhouette coefficient analysis makes it possible to select the optimal number of clusters by optimizing the separation distance between clusters. We normalized the gradients prior to clustering when calculations included all 8 gradients; if normalization is not performed, gradient 1 obscures the contribution of the last gradients given its much larger range of values.

### 2.5 Cerebello-cerebral functional connectivity (Supplementary methods)

We aimed to compare asymmetries between the two motor (IV/V/VI, VIII) and three nonmotor regions of cerebellar representation (VI/Crus I, Crus II/VIIB, IX/X) by comparing their relative position along diffusion embedding gradients. As a supplementary analysis, we also contrasted cerebello-cerebral connectivity from these regions using diffusion embedding gradient peaks within each of these areas of representation (e.g. contrasting cerebral cortical connectivity between first and second motor regions of representation). Cerebello-cerebral and intra-cerebellar connectivity Fisher’s z transformed values were obtained from the preprocessed HCP “dense connectome” (n=1003); maps were contrasted using the method for comparing correlated correlation coefficients described by Meng and colleagues^60^; and p maps were corrected at for multiple comparisons within the cerebral cortex using p<.05 voxel-based false discovery rate calculations.

## 3. RESULTS

### 3.1 Cerebellum gradients and relationship with discrete task activity and resting-state maps

Gradient 1 explained the largest part of variability in resting-state connectivity patterns within the cerebellum **(Fig. 1A)**. It extended bilaterally from lobules IV/V/VI and lobule VIII to posterior aspects of Crus I and Crus II as well as medial regions of lobule IX. Overlap with task activity maps **(Fig. 1B)** revealed that Gradient 1 is anchored at one end by cerebellar motor regions and at the other end by regions engaged in the language task of the HCP dataset. Regions situated between the two extreme ends corresponded to areas involved in working memory and emotion task processing. Social processing was diffusely distributed across Gradient 1. Overlap with cerebellar representations of cerebral cortical resting-state networks **(Fig. 1B)** revealed that Gradient 1 extends from sensorimotor network to DMN regions of the cerebellum. Ventral/dorsal attention and frontoparietal networks were situated between the two extreme ends.

**Fig. 1.**
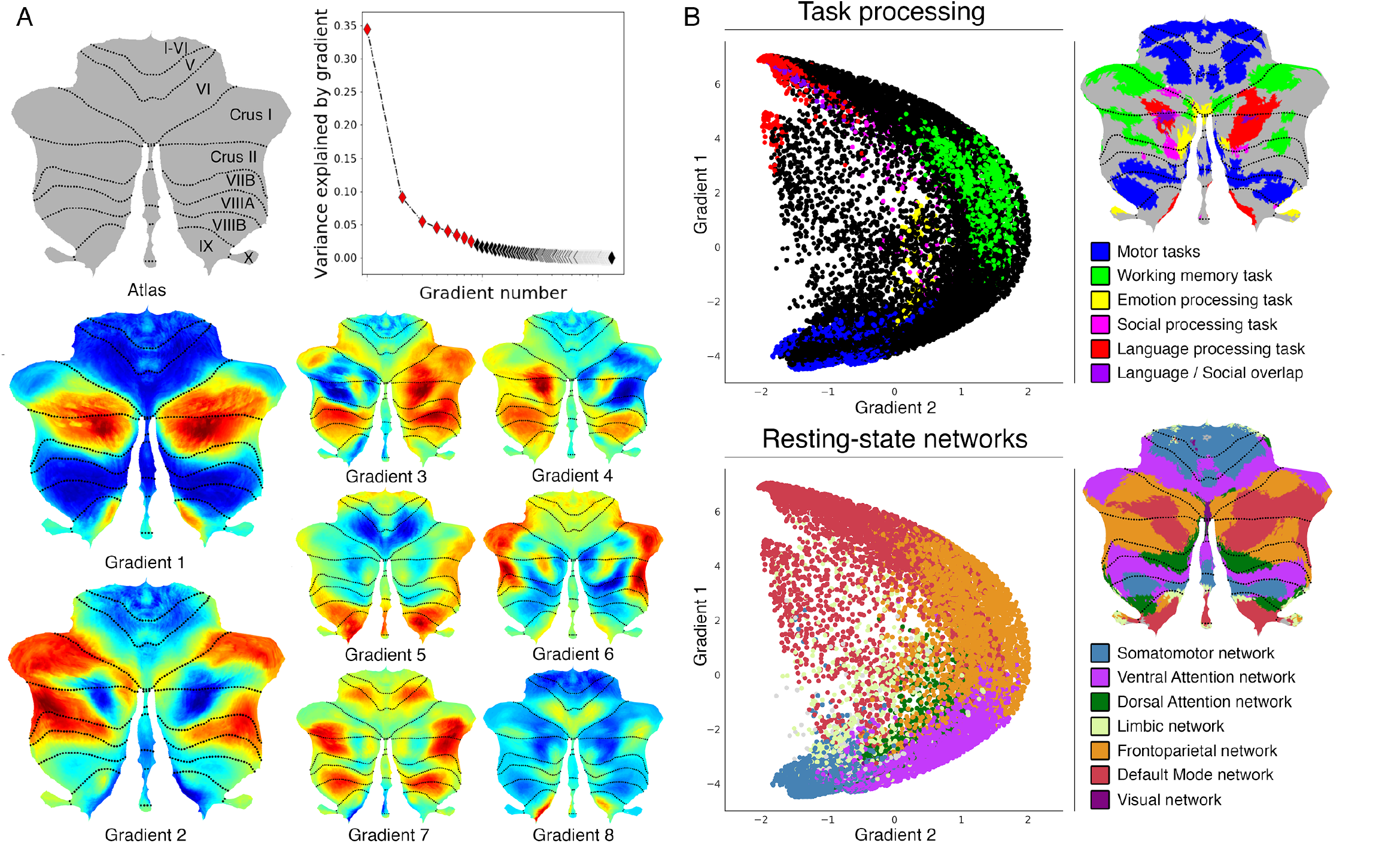
Cerebellum gradients and relationship with discrete task activity and resting-state maps. Gradient 1 extended from language task / DMN to motor regions. Gradient 2 isolated working memory / frontoparietal network areas. Gradients 3-4 revealed laterality asymmetries between nonmotor cerebellar regions. Gradients 5, 7 and 8 isolated aspects of lobule IX. **(A)** Top: Variance explained by each gradient; red diamonds correspond to the first eight gradients. Bottom: Gradients 1-8. **(B)** A scatterplot of the first two gradients. Each dot corresponds to a cerebellar voxel, position of each dot along x and y axis corresponds to position along Gradient 1 and Gradient 2 for that cerebellar voxel, and color of the dot corresponds to task activity (top) or resting-state network (bottom) associated with that particular voxel.

Gradient 2, the component accounting for the second-most variance, included at one end the anterior portions of Crus I and Crus II bilaterally **(Fig. 1A)**. These regions corresponded to areas engaged in the HCP working-memory task **(Fig. 1B)**. The same areas were included in cerebellar representations of the frontoparietal resting-state network. The other end of Gradient 2 included both regions involved in motor processing and regions involved in language processing; these areas correspond, respectively, to sensorimotor network and DMN regions.

Gradients 3 and 4 revealed laterality asymmetries in areas of nonmotor processing (Crus I, Crus II, VIIB and IX) but not in areas of motor processing (IV/V/VI and VIII) **(Fig. 1A)**. One extreme of Gradient 5, 7 and 8 included unilateral or bilateral aspects of lobule IX (**Fig. 1A**, extreme shown in red for Gradient 5 and 8 and in blue for Gradient 7). Gradient 6 included at one extreme anterior portions of Crus I and Crus II. The other end included medial Crus I and Crus II as well medial regions of lobule VIIB. The added percentage of variability explained by each following gradient was minimal.

Data from a single participant revealed a similar distribution of gradients 1 to 4 and a similar relationship with the same single subject motor, language and working-memory task processing **(Fig. S1)**.

**Fig. S1. (Supplementary figure).**
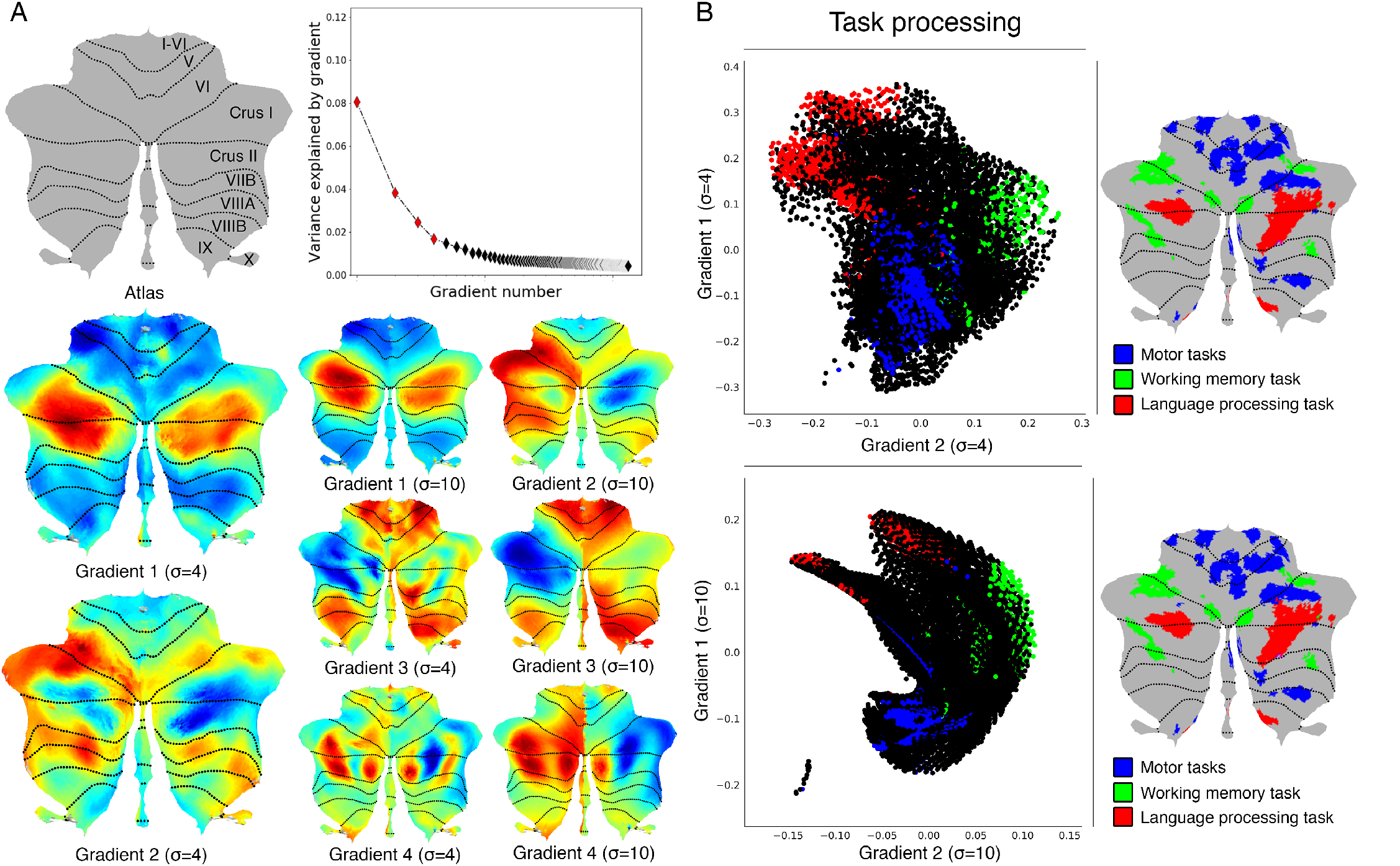
Cerebellum gradients and relationship with discrete task activity for an individual subject (one resting-state run of 15 minutes). **(A)** Top: Variance explained by each gradient; red diamonds correspond to the first four gradients. Bottom: Gradients 1-4 with 4 and 10 sigma (σ) values for smoothing of connectivity gradients. **(B)** A scatterplot of the first two gradients. Each dot corresponds to a cerebellar voxel, position of each dot along x and y axis corresponds to position along Gradient 1 and Gradient 2 for that cerebellar voxel, and color of the dot corresponds to task activity associated with that particular voxel. Scatterplots are shown for sigma = 4 (top) and sigma = 10 (bottom) values for smoothing of connectivity gradients. Task activity z maps are thresholded at z > 4 and include 4mm spatial smoothing as provided by HCP, in addition to sigma=2 smoothing using workbench command-cifti-smoothing.

Clustering of connectivity gradients revealed discrete networks similar to cerebello-cerebral connectivity parcellations from Buckner et al., 2011^36^ **(Fig. S2)**.

**Fig. S2 (Supplementary figure).**
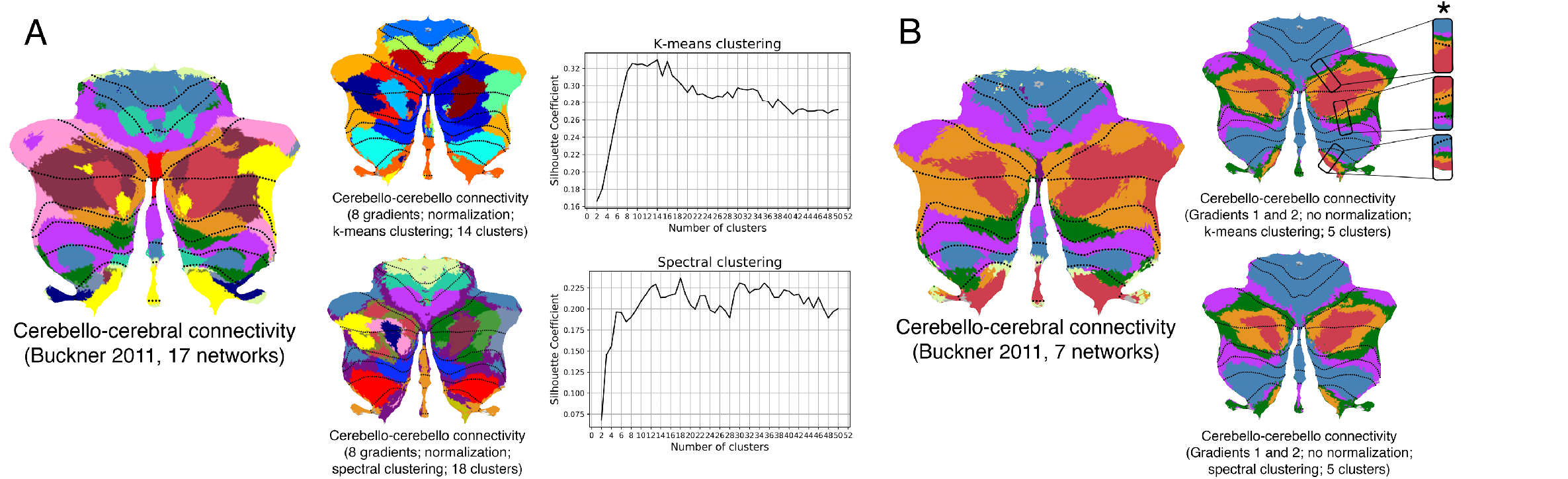
Clustering of connectivity gradients revealed discrete networks similar to cerebello-cerebral connectivity parcellations from Buckner et al., 2011^36^. **(A)** Silhouette coefficient analysis revealed an optimal number of 14 and 18 clusters for k-means and spectral cluster, respectively. Both clustering approaches revealed a distribution of parcels similar to the 17-network cerebello-cerebral connectivity parcellation from Buckner et al., 2011^36^. **(B)** K-means and spectral clustering using 5 clusters and gradients 1 and 2 revealed a distribution of parcels similar to the 7-network cerebello-cerebral connectivity parcellation from Buckner et al., 2011^36^, as well as a pattern of inverted representations (inverted pattern indicated with an asterisk). We used 5 clusters instead of 7 given the relatively small representation of two of the networks (visual and limbic) in Buckner et al., 2011, and the first two gradients given that those reflect the double/triple representation distribution which is observed in the 7-networks map from Buckner et al., 2011^36^. Note that while Buckner et al., 2011^36^ applied a winner-takes-all algorithm to determine the strongest functional correlation of each cerebellar voxel to one of the 7 or 17 cerebral cortical resting-state networks, our analysis included cerebellum-to-cerebellum correlations only.

### 3.2 Investigation of individual areas of motor and nonmotor representation

Resting-state as well as task processing analyses have revealed a cerebellar double motor (lobules IV/V/VI and VIII) and triple non-motor representation (lobules VI/Crus I, Crus II/VIIB and IX/X)^36,37^, but the functional significance of this distribution remains unknown. To investigate individual areas of motor and nonmotor representation, we isolated Gradient 1 highest 5% voxels within each area of nonmotor representation (“*High-G1*”), Gradient 2 highest 5% voxels within each area of nonmotor representation (“*High-G2*”), and Gradient 1 lowest 5% voxels within each area of motor representation (“*Low-G1*”) **(Fig. 2A)**. This parcellation is functionally meaningful because High-G1 / High-G2 correspond to different nonmotor task activity and resting-state network maps (language and DMN vs. working memory and frontoparietal/ventral-dorsal attention), and Low-G1 corresponds to areas of motor processing **(Fig. 1)**.

**Fig. 2.**
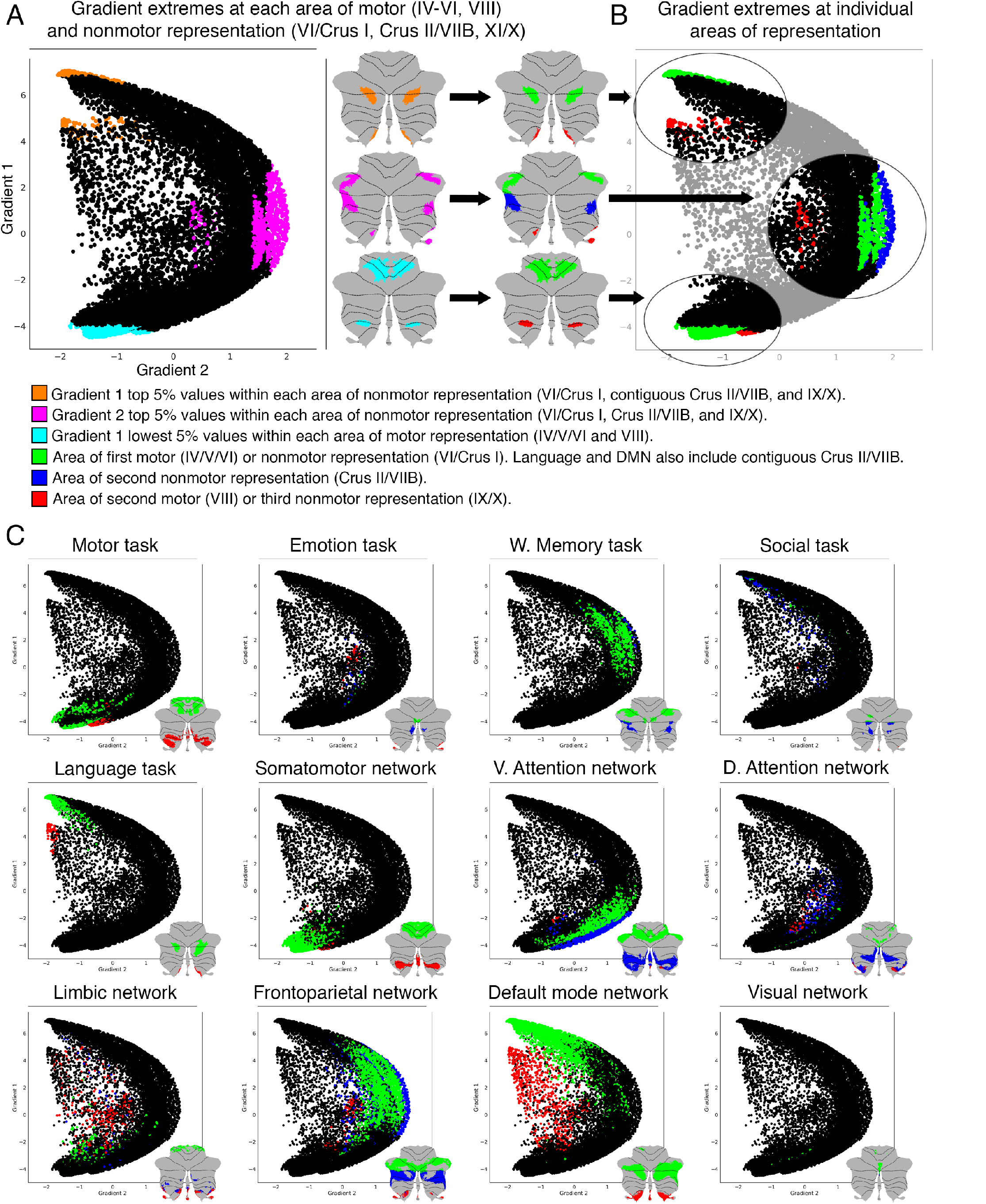
Investigation of individual areas of motor and nonmotor representation. Second motor and third nonmotor representations (shown in red) were consistently located at less extreme positions along Gradient 1 and/or 2. This hierarchical similarity might reflect a functional similarity - specifically, a less extreme level of information processing - between motor processing in lobule VIII and nonmotor processing in lobules IX/X. **(A)** Gradient 1 highest 5%, Gradient 2 highest 5%, and Gradient 1 lowest 5% voxels within each area of motor or nonmotor representation. **(B)** Plotting of the same areas shown in A, but differentiating each individual representation. **(C)** Plotting of all task activity and resting-state network maps differentiating each individual representation.

Further, we isolated each individual representation in task activity maps from Guell et al., 2018^37^ and resting-state network maps from Buckner et al., 2011^36^. Specifically, we isolated first motor (VI/V/VI) and second motor representation (VIII) of the motor task map and somatomotor network. Language task and DMN were separated in first and contiguous second nonmotor (VI/Crus I/Crus II/VIIB) and third nonmotor (IX/X) representations, given the contiguous first and second representations of these maps in Crus I/Crus II. All other tasks and resting-state maps were divided in first nonmotor (VI/Crus I), second nonmotor (Crus I/VIIB) and third nonmotor representation (IX/X).

When analyzing the position of each individual representation along Gradient 1 and 2, second motor and third nonmotor representations were consistently located at less extreme positions along these two gradients. We observed this consistently across gradient-derived parcellations (High-G1/Hig-G2/Low-G1, **Fig. 2B**) as well as task and resting-state network maps **(Fig. 2C)**. Second motor and third nonmotor representations are shown in red in **Fig. 2B** and **Fig. 2C**. Note, for example, that second motor representation in Low-G1, motor task and somatomotor network was located at a less extreme position along Gradient 2. Following a similar logic, third representation of High-G1, language task and DMN was located at a less extreme position along Gradient 1. Third representation of High-G2 and frontoparietal network was located at a less extreme position along Gradient 2. Third representation of emotion, social task, ventral attention, dorsal attention and limbic networks showed a less clear distribution, but was nonetheless consistently located at more central (i.e., less extreme) positions along Gradient 1 and/or 2. This organization could not be observed in working memory task and visual network given that these maps were not represented in lobules IX/X.

A data-driven clustering of the first two gradients resulted in a division of gradients 1 and 2 in three areas encompassing our High-G1/High-G2/Low-G1 parcellation **(Fig. S3)**, further supporting this hypothesis-driven division. Crucially, the same relationship between the two motor and three nonmotor areas of representation was observed in the analysis of a single subject with only one resting-state run of 15 minutes **(Fig.S4)**. A supplementary cerebello-cerebral connectivity analysis revealed additional differences in cerebral cortical connectivity from each area of representation **(Fig. S5)**.

**Fig. S3 (Supplementary figure).**
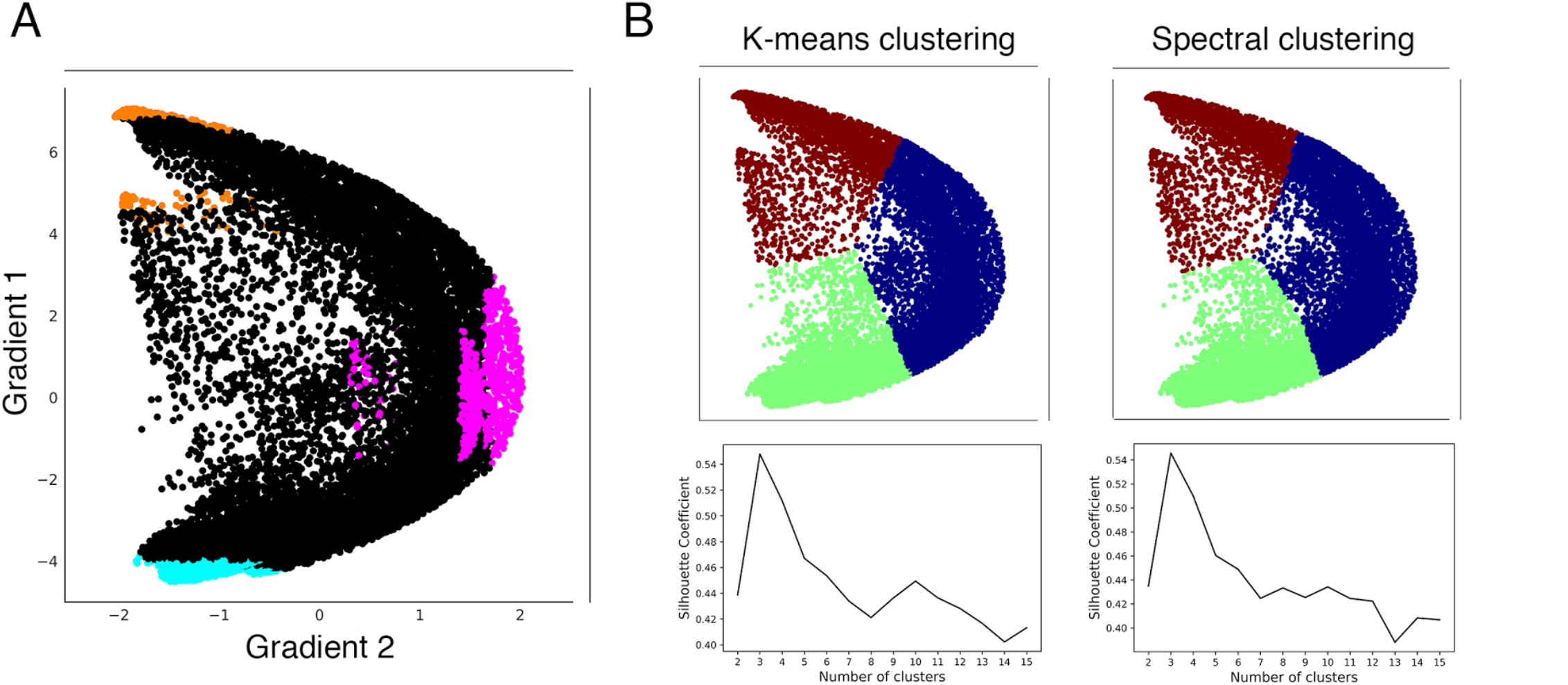
**(A)** Our hypothesis-driven parcellation based on Gradient 1 lowest 5% voxels at each area of motor representation (blue), and Gradient 1 (orange) and Gradient 2 (pink) highest 5% voxels at each area of nonmotor representation. **(B)** Using gradients 1 and 2 after normalization, Silhouette Coefficient analysis of k-means and spectral clustering reveals that three is the optimal number of clusters. K-means and spectral clustering using three clusters reveals a separation that encompasses our hypothesis-driven High-G1/Hig-G2/Low-G1 division.

**Fig. S4 (Supplementary figure).**
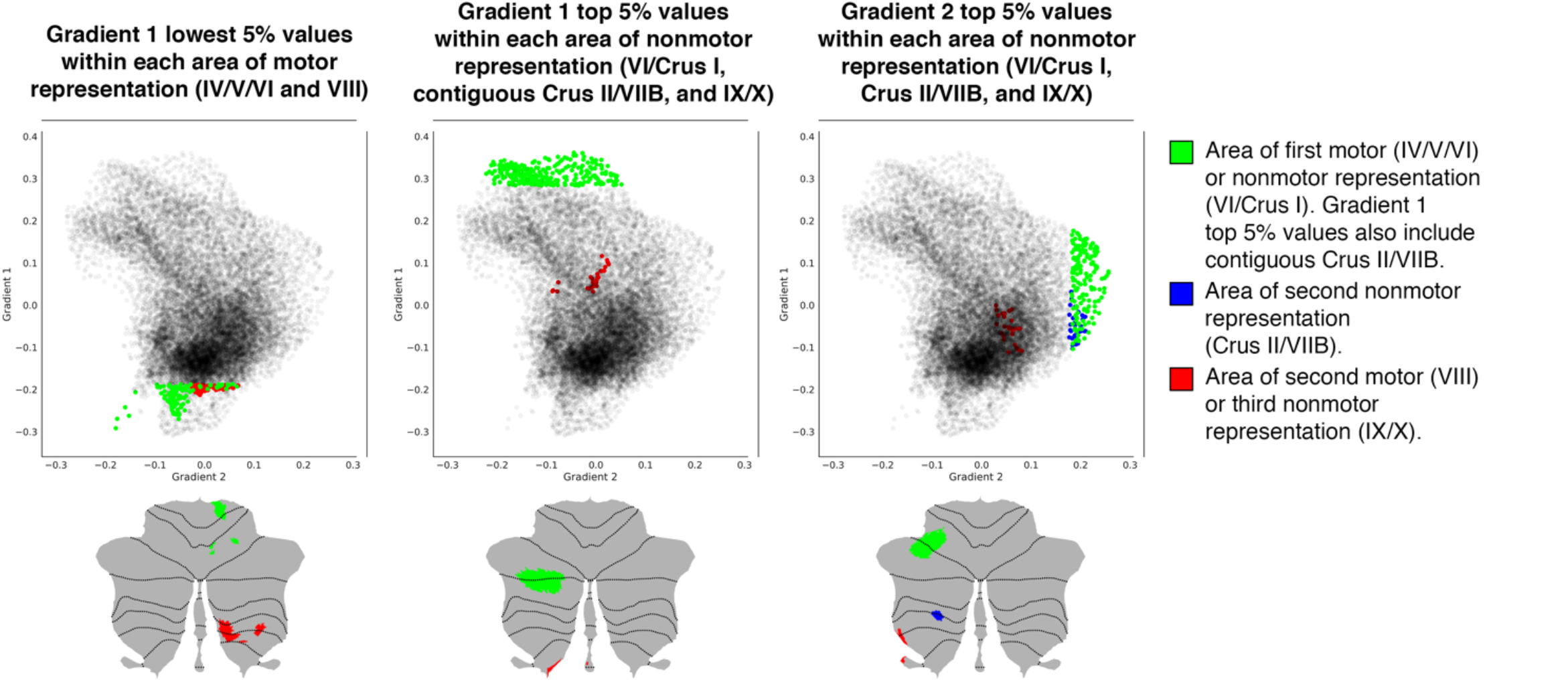
Investigation of individual areas of motor and nonmotor representation using data from one single subject (one resting-state run of 15 minutes). As in the group analysis, second motor and third nonmotor representations (shown in red) were consistently located at less extreme positions along Gradient 1 and/or 2. Given the asymmetries between left and right hemisphere in the analysis of a single subject (see Fig.S1), maps in these figures include only the right hemisphere for Gradient 1 lowest 5% values and only the left hemisphere for Gradient 1 and 2 highest 5% values. Opacity of black dots in these plots was decreased to improve the visibility of green, blue and red dots. Gradients were smoothed with sigma=4.

**Fig. S5 (Supplementary figure).**
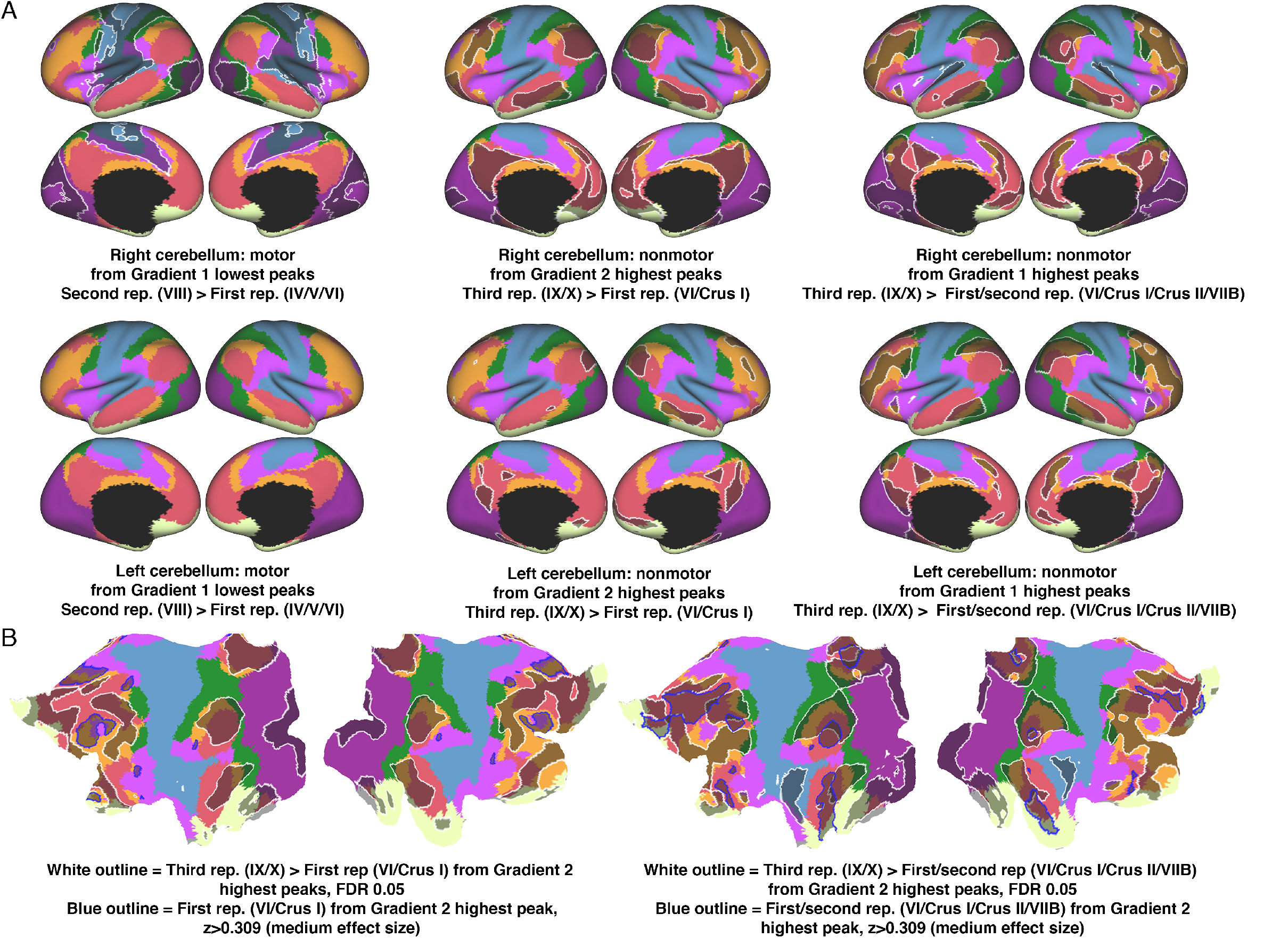
Contrasts of cerebello-cerebral connectivity from Gradient 1 or 2 peaks at each area of motor or nonmotor representation. Lowest values of Gradient 1 target motor regions. Lowest value of Gradient 1 in the area of second motor representation revealed stronger connectivity with areas close to, but not at, primary motor and somatosensory cortex **(A, left column)**. Higher values of Gradient 1 target DMN regions **(B, right image)**. Gradient 1 peak at the area of third nonmotor representation revealed stronger connectivity with DMN/frontoparietal regions **(A, right column)**. Higher values of Gradient 2 target ventral attention network regions **(B, left image)**. Gradient 2 peak at the area of third nonmotor representation revealed stronger connectivity with DMN/frontoparietal regions **(A, middle column)**. We interpret all these connectivity contrasts as a reflection of a less extreme level of information processing along the motor / nonmotor dimension (from primary motor/somatosensory cortex [maximum motor] to regions surrounding these structures, and from DMN [maximum non-motor] to frontoparietal/DMN areas); or along the task-focus / task-unfocused dimension (from ventral attention [maximum task-focus] to frontoparietal/DMN areas).

## 4. DISCUSSION

This is the first study to investigate the progressive, hierarchical organization of the cerebellum. Contrasting with the fundamental and well-established primary-unimodal-transmodal hierarchical organization in the cerebral cortex^1,2^, the principal axis of cerebellar motor and nonmotor organization remains unknown. We describe for the first time that cerebellar functional regions follow a gradual organization which progresses from primary (motor) to transmodal (DMN, task-unfocused) regions. Further, the relationship between the two principal gradients and the two motor and three nonmotor areas of representation revealed hierarchical similarities – perhaps reflecting functional similarities – between nonmotor processing in lobules IX/X (third nonmotor representation) and motor processing in lobule VIII (second motor representation). These interpretations are further supported by data-driven clustering and cerebello-cerebral functional connectivity analyses. Importantly, these descriptions remain observable at an individual level. These findings, from an exceptionally large and high-quality dataset, provide new and fundamental insights into the functional organization of the human cerebellum, unmask new testable questions for future studies, and yield an unprecedented tool for the topographical interpretation of cerebellar findings.

### 4.1 Gradient 1 extends from motor to nonmotor areas: cerebellar macroscale organization is sensorimotor-fugal

Gradient 1 extended from regions corresponding to motor task activity and sensorimotor network representation to regions corresponding to language task activity and DMN representation **(Fig. 1B)**. The overlap between language task activity and DMN may be due to the language task contrast which subtracted listening to stories minus answering arithmetic questions. This subtraction may capture processes similar to those that engage DMN regions, such as autobiographical memory retrieval, daydreaming and conceiving the perspective of others^61^. Consistent with this hypothesis, HCP language task activity also overlapped with DMN in the cerebral cortex **(Fig. S6)**. Working memory task processing was situated at a middle point along Gradient 1, similar to the distribution of frontoparietal and ventral attention networks **(Fig. 1B)**. It is reasonable to conceptualize working memory as a nonmotor task which is not as distant from motor function as a story listening task, justifying its middle position along Gradient 1. Similarly, tasks that activate DMN regions such as daydreaming and mind wandering^62–64^ can be conceptualized as more distant from motor processing than goal-directed cognitive control and decision-making processes that activate frontoparietal network regions^65^. Ventral and dorsal attention networks were located far from DMN along Gradient 1 **(Fig. 1B)**, consistent with the view that DMN and ventral/dorsal attention networks are two opposing brain systems^66^. The frontoparietal network is conceptualized as a mediator between the two^65^, justifying its position between ventral/dorsal attention networks and DMN along Gradient 1 **(Fig. 1B)**. This conceptualization of Gradient 1 is also coherent with a previous report analyzing cerebellar activity at multiple time points, from motor planning to motor output^67^. The authors described a lateromedial succession *“from will to action”* (see Figure 3 in Hulsmann et al., 2003^67^) which is in accordance with the direction of Gradient 1 from nonmotor to motor regions in our analysis.

This is the first study to report a sensorimotor-fugal macroscale organization in the cerebellum; i.e., a hierarchical organization that progresses away from sensorimotor function. A different study using diffusion map embedding analysis in the cerebral cortex reported similar results^34^. In that case, the principal gradient extended from primary cortices (visual, somatosensory/motor and auditory) to regions corresponding to the DMN. As in the cerebellum, the frontoparietal network was also located between DMN and ventral/dorsal attention networks, and working memory task activity was also located at a middle position along the principal gradient. Of note, the cerebellum does not show functional connectivity with primary visual or auditory cortices^36^, but is anatomically and functionally connected with areas of primary sensorimotor processing and consistently engaged in simple motor tasks. It is therefore reasonable to consider that a gradient from motor to DMN areas in the cerebellum is the equivalent of a gradient from motor/visual/auditory to DMN areas in the cerebral cortex.

This finding strongly suggests that cerebellum and cerebral cortex share a similar macroscale principle of organization, namely, that both structures share a hierarchical organization which gradually progresses away from unimodal streams of information processing. While this organization has long been defended in the cerebral cortex^1,34,35^, the present analysis is the first to reveal an analogous principle in the cerebellum. This is a notable observation because of two reasons. First, gradients obtained in our analysis correspond to intrinsic connectivity profiles of cerebellar voxels with the rest of the cerebellum only, rather than with the rest of the brain. Therefore, our analysis reflects the organization of the cerebellum without invoking its connectivity profiles with the cerebral hemispheres. In this way, the fact that we observed a similar principle of organization between the cerebral cortex and the cerebellum does not constitute an imposition of our method of analysis (unlike in Buckner et al., 2011^36^). Second, the cerebral cortical notion that there is a hierarchical organization which gradually progresses away from unimodal streams of information^1,34,35^ is implicitly predicated on the anatomical knowledge that there are synapses linking adjacent cerebral cortical regions. However, no cortico-cortical anatomical connections exist in the cerebellum. It is therefore nontrivial to observe this parallel organization in the cerebellum, the anatomical origin of which may be addressed in future studies. We speculate that such a functional organization could be a consequence of the arrangement of cerebello-cerebral anatomical connections which might affect correlations in resting-state activity between cerebellar regions. The same possibility raises further questions regarding the precise distribution of cerebello-cerebral anatomical connections that would be required to achieve such a parallel mapping of functional gradients in the cerebral cortex and cerebellum.

The finding that a similar distribution of the first two gradients and their relationship with motor, language and working memory task processing can be observed at an individual level **(Fig. S1)** supports the assertion that this organization is not an artifact generated as a result of averaging a large number of subjects, and highlights the potential application of this fundamental principle in future small group or single subject investigations.

**Fig. S6.**
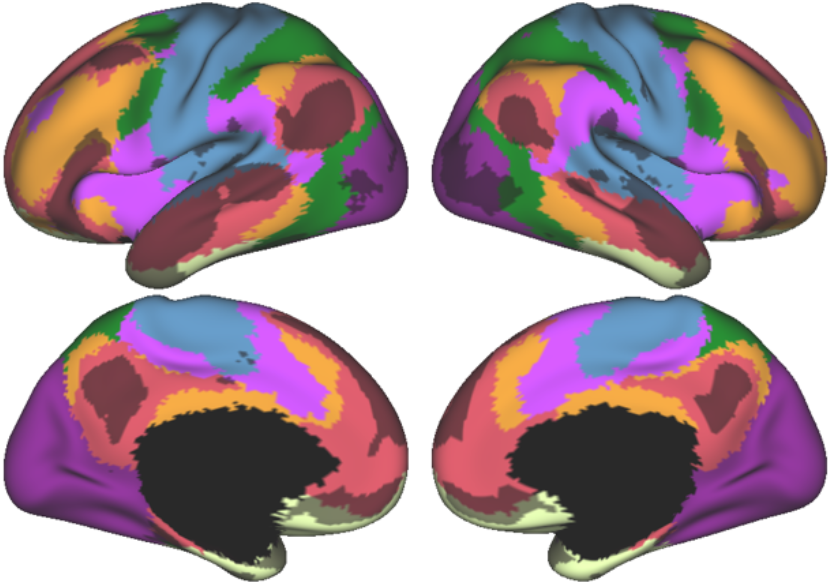
Cerebral cortical resting-state networks from Yeo and colleagues^33^ revealed an overlap between DMN (red) and language task activity (grey) also in the cerebral cortex. Language task activity map corresponds to Cohen’s d map thresholded at 0.5 from Guell et al., 2018^37^.

### 4.2 Integrating Gradient 1 and Gradient 2: task processing in the cerebellum understood in terms of distance from motor processing and amount of task-focus

Gradient 1 extended from motor to nonmotor (task-unfocused, DMN) regions. In contrast, Gradient 2 (the component accounting for the second-most variance) was anchored at one end by working memory task and frontoparietal network regions. The other extreme of Gradient 2 corresponded to both extremes of Gradient 1, namely, (i) regions corresponding to motor task activity and sensorimotor network and (ii) regions corresponding to language task activity and DMN representation. The functional significance of this distribution might be analyzed as follows. Working memory HCP task corresponds to *Two back* (respond if current stimulus matches the item two back) minus *Zero back* (respond if current stimulus matches target cue presented at start of block). HCP language task correspond to *Story* (listen to stories) minus *Math* (answer arithmetic questions). HCP motor tasks corresponds to *Movement* (tap left fingers, or tap right fingers, or squeeze right toes, or squeeze left toes, or move tongue) minus *Average* (average of the other four movements). What dimension corresponding to the working memory task is equally absent in the language and the motor task contrasts? One possible explanation is task focus. Whereas the working memory task contrast isolates a higher load of working memory (therefore a higher load of task focus), task focus is eliminated from the language task contrast after subtracting the math condition, and task focus is eliminated from the motor task contrast after subtracting the average of other movements. Coherently, frontoparietal and ventral attention networks (the extreme of Gradient 2, **Fig.1B**) are task-positive networks^65^ while DMN and somatosensory network (the other extreme of Gradient 2) are not.

In this way, Gradient 1 and Gradient 2 classify information processing in the cerebellum along two dimensions: distance from motor processing (Gradient 1) and amount of task-focus (Gradient 2). HCP motor task contrast isolates pure motor processing and eliminates task-focus demands. In consequence, HCP motor task is situated at a minimal position in Gradient 1 (i.e., maximally motor) and at a minimal position in Gradient 2 (i.e., minimally task-focused) **(Fig. 1B)**. HCP working memory task isolates a higher load of working memory by subtracting a two-back minus a zero-back condition. The isolated cognitive process is closely related to task focus and is therefore situated at a maximum position in Gradient 2. At the same time, working memory represents a nonmotor process and is therefore situated higher than the HCP motor task along Gradient 1. This notwithstanding, working memory is situated lower than the HCP language task along Gradient 1. This order seems logical by considering that goal-nondirected processes targeted by the HCP language task contrast are more distant from pure motor processing than those goal-directed processes isolated by the working memory task contrast. Similarly, mind-wandering states are, by definition, task-unfocused, explaining the position of the HCP language task at the lowest extreme of Gradient 2.

Our interpretation of task focus in the cerebellum in terms of distance from motor processing and amount of task-focus is also coherent with the general distribution of data points when plotting Gradient 1 against Gradient 2 (see plots in **Fig. 1B**). First, there are no cerebellar voxels with simultaneous maximum Gradient 1 and Gradient 2 values. Maximum Gradient 1 values correspond to DMN regions, and DMN processes are task-unfocused by definition. Therefore, Gradient 1 maximum values must have low Gradient 2 values. Second, there are no cerebellar voxels with simultaneous minimum Gradient 1 and maximum Gradient 2 values. This distribution is consistent with the notion that increasing attentional demands of a motor task adds nonmotor computational demands. Accordingly, Gradient 1 lowest values cannot increase their position along Gradient 2 without simultaneously acquiring a higher position along Gradient 1.

While HCP motor, working memory and language task activity maps were situated at extreme regions along Gradient 1 and/or 2 **(Fig. 1B)**, social and emotion processing did not adhere to any extreme along these gradients. This observation may provide novel insights into the organization and nature of social and emotion processing in the cerebellum. Social processing task activity map spanned across Gradient 1, perhaps reflecting a multimodal nature of social processing in the cerebellum in the dimension of motor to nonmotor processing. The conceptualization of social processing in the cerebellum as an activity that engages multiple levels of information processing along the motor-nonmotor dimension may relate to the concomitant impairment of social skills, nonmotor tests such as Rey’s figure or Tower test, and some motor abilities (e.g. equilibrium and limb coordination) in autism spectrum disorders^68^.

Emotion processing was situated at a central position in both Gradient 1 and 2. We understand this distribution as an inability to clearly classify emotion processing along the gradients of distance from motor processing (Gradient 1) and amount of task focus (Gradient 2). Higher working memory load as isolated by the HCP working memory task corresponds to a level of information processing with high task-focus demands. At the same time, the subtraction of the HCP language task isolates task-unfocused processes which are maximally removed from pure motor processing. The results of the HCP emotion processing task contrast, on the other hand, are not as well defined along these dimensions. The subtraction of *Faces* (“decide which of two angry/fearful faces on the bottom of the screen match the face at the top of the screen”) minus *Shapes* (same task performed with shapes instead of faces) isolates higher emotional content in the information that is processed. It may be argued that this higher emotional content corresponds to an intermediate position between pure motor and high-nonmotor level information processing (explaining the intermediate position along Gradient 1), and that this higher emotional content results in mildly increased task focus (explaining the intermediate position along Gradient 2).

### 4.3 Confirmation of the double/triple representation hypothesis

Resting-state as well as task processing analyses have revealed a cerebellar double motor (lobules IV/V/VI and VIII) and triple non-motor representation (lobules VI/Crus I, Crus II/VIIB and IX/X)^36,37^. The distribution of Gradient 1, the component that explains the greatest variability in resting-state intra-cerebellar connectivity patterns, confirms this organization. Gradient 1 lowest values correspond to lobules IV/V/VI and VIII (**Fig. 1A**, dark blue regions in Gradient 1), demarcating the two areas of motor representation. The highest values correspond to lobules Crus I, Crus II, and lobule IX (**Fig. 1A**, dark red regions in Gradient 1) - these regions correspond to the first, contiguous second, and third nonmotor representation areas, respectively. Taken together, the double motor / triple nonmotor organization has now been shown in cerebellar representations of cerebral resting-state networks^36^, cerebellar task activity^37^, cerebro-cerebellar functional connectivity from cerebral cortical task activity peaks^37^, and gradients of intra-cerebellar patterns of functional connectivity (the present study). Gradient 2 also revealed a similar distribution, with its maximum values located in Crus I, Crus II/VIIB, and lobules IX/X.

Clustering of connectivity gradients revealed discrete networks resembling cerebello-cerebral connectivity parcellations in Buckner et al., 2011^36^, and replicating their double/triple representation distribution **(Fig. S2)**. This observation supports the generalizability of the double/triple representation hypothesis to multiple directions of functional connectivity, namely, cerebello-cerebral and intra-cerebellar.

A *“network approach to the localization of complex functions”* rather than *“an exclusive concentration of function within individual centers in the brain”^69^* has long been adopted in the cerebral cortex^33,69-71^, although some complex functions are indeed organized into focally specific brain regions^72,73^. Accumulating evidence for a double motor / triple nonmotor organization in the cerebellum warrants an analogous shift in the understanding of cerebellar functional neuroanatomy. Just as *“each distributed network consists of association areas spanning frontal, parietal, temporal and cingulate cortices”^69^,* the data indicate that each nonmotor cerebellar network consists of three representations spanning VI/Crus I, Crus II/VIIB and IX/X. There are no intrinsic anatomical connections linking these cerebellar areas, but tract tracing studies in monkeys hint at the possibility of an anatomical correlate of the double motor / triple nonmotor organization. This conclusion is based on shared cerebello-cerebral cortical loops: lobules VI/V/VI and VIII receive input from and project to M1, and lobules Crus I/Crus II and IX/X receive input from and project to area 46^76^. Further, as in the cerebral cortex, distributed networks may exist adjacent to each other within each area of nonmotor representation in the cerebellum. In the same way that *“adjacent areas in the parietal cortex belonging to separate networks are differentially connected to adjacent areas of corresponding networks in the frontal, temporal and cingulate cortices”* ^33,74-76^, adjacent areas in VI/Crus I belonging to separate networks are differentially related to adjacent areas of corresponding networks in Crus II/VIIB and IX/X. This is revealed by non-overlapping nonmotor task activity maps within each area of representation in Guell et al., 2018^37^, the unmasking of multiple resting-state networks within each area of representation in Buckner et al., 2011^36^, and the distribution of Gradient 1 in the present analysis. Task contrasts or connectivity analyses might reveal incomplete engagement of the triple nonmotor cerebellar network - a discussion regarding this incomplete engagement would be appropriate in these cases. For instance, incomplete engagement of the triple nonmotor network might be functionally meaningful, e.g., activity in the areas of first and second representations, but not in the area of third representation. Similarly, future studies may discuss group contrasts where a given neurological or psychiatric disease results in functional or structural cerebellar abnormalities within only one area of motor or nonmotor representation. This approach might be critical for the understanding of cerebellar systems physiology and pathophysiology. Consequently, a critical next step towards a more comprehensive and nuanced understanding of cerebellar functional neuroanatomy is the investigation of distinct contributions of each area of motor and nonmotor representation. The following section addresses this question.

### 4.4 Second motor (VIII) and third nonmotor representation regions (IX/X) are situated at a less extreme level along Gradient 1 and/or 2: hierarchical similarities between lobules VIII and IX/X suggest functional similarities between third nonmotor and second motor representation

A review^77^ frames the question of “the functional significance of the two (or three) cortical representation maps in the cerebellum” as one of the principal outstanding enigmas in cerebellar neuroscience. Our present study provides the data to attempt to address this question for the first time, as follows.

Second motor representation (lobule VIII) and third nonmotor representations (lobule IX/X) were consistently located at less extreme positions along Gradient 1 and/or 2 when compared to their first motor and first/second nonmotor representations, respectively. This pattern was consistently observed in all maps analyzed, including gradient-derived cerebellar parcellations **(Fig. 2B)**, task activity maps (from Guell et al., 2018^37^) and resting-state maps (from Buckner et al., 2011^36^) **(Fig. 2C)**. Further, this distribution was also observed in 15 minutes of resting-state data in a single subject **(Fig. S4)**. This observation suggests that the nonmotor contribution of lobules IX/X (third nonmotor representation) is different from the nonmotor contribution of lobules VI/Crus I/Crus II/VIIB (first and second nonmotor representations), and that the motor contribution in lobule VIII (second motor representation) is different from the motor contribution of lobules IV/V/VI (first motor representation). This conservative conclusion is, on its own, novel in the field of cerebellar systems neuroscience. We further speculate that a less extreme position along Gradient 1 and/or 2 in both third nonmotor and second motor representation represents, in both cases, a less extreme level of information processing. “Extreme” here refers to the poles of the sensorimotor-fugal organization (Gradient 1) and the task-focus/task-unfocus organization (Gradient 2). Specifically, a less extreme position along Gradient 1 corresponds to a less extreme level of information processing along the motor / nonmotor dimension, and a less extreme position along Gradient 2 corresponds to a less extreme level of information processing along the task-unfocused / task-focused dimension. Because this pattern of a less extreme level of information processing is observed in both second motor representation and third nonmotor representation, we argue that nonmotor activity in lobules IX and X (third nonmotor representation) might emerge from, and follow the logic of, motor processing in lobule VIII (second motor representation). This notion is inspired by the organization of the cerebral cortex where multimodal or association cortical areas are related to their nearby unimodal areas. For example, Broca’s area is close to the primary motor cortex, while Wernicke’s area is close to the primary auditory cortex. The analogy in the cerebellum is that nonmotor activity in lobules IX and X is adjacent to, and therefore follows the logic of, motor activity in lobule VIII. Restated, the relationship between first motor and second motor representation resembles the relationship between first/second nonmotor and third nonmotor representations, just as the relationship between primary motor and primary auditory cortex reflects the relationship between Broca’s area and Wernicke’s area.

The data show that the second representation of motor task activity, sensorimotor network and “Low-G1” (motor) maps were consistently located at a higher position along Gradient 2 when compared to their first representation. This suggests that while the first motor representation is engaged in pure motor processing as isolated by the *Movement* (e.g. tap left fingers) minus *Average* (average of the other four movements) contrast, second motor representation is engaged in motor process that require higher task focus. In this way, second motor representation corresponds to a less extreme level of task-unfocused motor information processing. Following a similar logic, third representation of language task, DMN and “High-G1” maps were consistently located at a lower position along Gradient 1 when compared to their first and contiguous second representations. While these first and contiguous second representations are at an extreme level of information processing (i.e., maximally nonmotor), third representation is in a less extreme position (i.e., less extreme in the motor/nonmotor dimension). Also consistent with this logic, the third representation of working memory, frontoparietal network and “High-G2” maps were consistently located at a lower position along Gradient 2 when compared to their first and second representations. These first and second representations were at an extreme level of information processing – specifically, maximally task-focused. The third representation was located further from this extreme, i.e., less extreme in the task-unfocused/task-focused dimension. Ventral and dorsal attention networks were not located at one clear gradient extreme, but their distribution of three representations also followed the logic that third representation (lobule IX/X) was located at a less extreme position along Gradient 1 and/or 2.

Of note, the second representation of working memory, frontoparietal network and “High-G2” was located similar to its first representation. This proximity between the first and second representations indicates that the relationship between second motor and third nonmotor representation does not apply to the relationship between second motor and second nonmotor representation. Restated, nonmotor processes in lobules IX/X share hierarchical principles with motor processing in lobule VIII (an analogous “less extreme” level of information processing) - in contrast, this relationship does not apply between nonmotor processing in lobules Crus II/VIIB and motor processing in lobule VIII.

A cerebello-cerebral connectivity analysis further supports the hypothesis that second motor and third nonmotor regions of representation correspond to a less extreme level of information processing when compared to their first motor and first/second nonmotor representations, respectively **(Fig. S5)**.

The constellation of symptoms that follow cerebellar strokes of the posterior inferior cerebellar artery (PICA) may also support our hypothesis. PICA occlusion commonly results in the infarction of lobule VIII (second motor representation) but not of lobules IV/V/VI (first motor representation). Notably, these lesions result in little or no motor deficits^78,79^. Our hypothesis that second motor representation corresponds to a less extreme level of pure motor information processing might explain the lack of pure cerebellar motor symptoms (gait ataxia, appendicular dysmetria, dysarthria) after PICA stroke. Whereas the pattern of deficits arising from lesions of the second motor representation may go undetected with the standard neurological motor examination, our data predict that fine discriminative testing may reveal deficits in motor-related tasks that require high task focus. This might include motor performance abnormalities that only manifest in the presence of distractors. However, PICA strokes also damage other lobules such as Crus II and VIIB - deficits in motor tasks requiring high task focus may be difficult to dissociate from nonmotor abnormalities arising from infarction of cerebellar regions other than lobule VIII. We are not aware of any report of isolated lobule VIII injury in humans - however, Dow, 1938^80^ performed isolated ablation of lobule VIII in three rhesus monkeys. The author reported that *“In all 3 animals in which the pyramis* (i.e. lobule VIII) *alone was damaged little that was abnormal could be detected, except that the animal when running down a long corridor apparently was unable to stop quickly enough to avoid crashing head-on against the end wall. No visual defect was present. The abnormality was never observed later than the third or fourth day after operation”.* Aberrant motor behavior in the absence of classical cerebellar motor symptoms may be consistent with our reasoning. fMRI task activity analyses have made claims regarding distinct functional contributions of the cerebellar second motor representation^81–85^; however, none has demonstrated statistically significant lobule VIII activity in the absence of lobule IV/V/VI activity for any given task contrast. Kipping and colleagues^86^ reported lobule VIII functional connectivity with cerebral cortical regions other than motor and premotor regions, a pattern of connectivity consistent with our hypothesis that the second motor representation is located at a less extreme level of motor processing.

We showed that during a working memory task there was activity in the cerebellum in the first and second nonmotor representations, but not in the third representation^37^. In contrast, functional connectivity was observed in all three areas of representation when seeding from the cerebral cortical peak of the working memory task. In the light of the present observations, our interpretation is that functional connectivity revealed the full pattern of triple representation of task-focused mid-nonmotor processing areas, but when engaged with a working memory task, the third representation in the network was not recruited due to excessive task-focus demands (i.e., due to an extreme level of information processing along the task-unfocused/task-focused dimension).

Some anatomical peculiarities of lobules IX/X conform to the notion of a functionally distinct nonmotor contribution of these lobules. Glickstein and colleagues^87^ reported that the principal target of pontine visual cells in monkeys is lobule IX. A specific type of cell, the Calretinin-positive unipolar brush cell, is preferentially located in lobules IX and X in many species^88,89^ and receives vestibular afferents^90^. Accordingly, lobules IX and X are classically considered to represent the vestibulocerebellum. One highly speculative proposal is that the incorporation of visual/vestibular streams of information in lobules IX/X, but not in lobules VI/Crus I/Crus II/VIIB, might be related to the asymmetries we describe between the third and the first/second nonmotor representations. Indeed, some lines of study investigate the link between vestibular function and limbic and cognitive functions including visuospatial reasoning^91–93^. The notion that asymmetry between nonmotor representations may arise from heterogeneity in cerebellar patterns of connectivity, rather than cytoarchitecture or physiology, is in accord with the notion of a Universal Cerebellar Transform^5,94,95^.

### 4.5 Additional gradients

Gradients 3 and 4 revealed asymmetries between left and right hemispheres. Notably, these asymmetries were constrained to areas of nonmotor processing (see **Fig. 1A**, asymmetries are not present in motor lobules IV/V/VI and VIII). Consistent with this observation, left/right asymmetries in cerebellar processing have been described in multiple nonmotor tasks^96^ but not in motor processing. Gradient 5, 7 and 8 isolate aspects of lobule IX at one extreme of their distribution **(Fig. 1A)**. A separation of lobule IX (the area of the third area of nonmotor representation) is harmonious with the present observation that nonmotor processing in lobules IX and X is distinct from nonmotor processing in lobules VI, Crus I, Crus II and VIIB. Gradient 6 extended from anterior aspects of Crus I, Crus II and VIIB to medial regions of these lobules. Task activity overlap with this gradient did not reveal a clear interpretable pattern, perhaps representing additional principles of organization that may be unmasked by future studies of cerebellar connectivity gradients.

### 4.6 Relevance for future investigations

This is the first study to describe the principal gradient of macroscale function in the cerebellum. Following a logic similar to the fundamental and well-established primary-unimodal-transmodal hierarchical organization in the cerebral cortex^1,34^, we report that cerebellar macroscale organization is sensorimotor-fugal. Regions further from the central aspect of lobules IV/V/VI and VIII are, accordingly, further from cerebellar motor function in a gradient from motor to maximally non-motor (mind-wandering, non-goal oriented) function. This concept is analogous to the well-established knowledge in the cerebral cortex that regions progressively further from primary cortices (motor/somatosensory, auditory, visual) are progressively involved in more abstract, transmodal, non-primary processing. This fundamental concept has greatly influenced topographical investigations in the cerebral cortex, and it is reasonable to consider that the present description may equally influence cerebellar investigations. The publicly available cerebellum gradient maps from the present study in multiple file formats and structural spaces (https://github.com/gablab/cerebellum_gradients, folder “FINAL_GRADIENTS”) will facilitate the inclusion of the sensorimotor-fugal principle of cerebellar macroscale organization in future investigations.

The distribution of these principal gradients confirmed the double motor / triple nonmotor organization in the cerebellum, highlighting the need to refer to this organization when discussing cerebellar functional or structural findings. Close attention to this network organization may become critical for the understanding of cerebellar structure and function in health and disease. Clusters in lobules IV/V/VI and VIII are commonly interpreted as first and second representations of motor processing. The same reasoning should be applied to nonmotor findings, for example, in the interpretation of degeneration of Crus I and IX in Alzheimer’s disease^27^.

One important secondary implication of the analysis of connectivity gradients in the present study is the unmasking of hierarchical similarities between second motor (VIII) and third nonmotor (IX/X) representations in gradient-derived parcellations, task activity and resting-state maps. We interpret this relationship as an indication that nonmotor processing in lobules IX/X emerges from, and follows the logic of, motor processing in lobule VIII – specifically, processing in both regions corresponds to a less extreme level of information processing when compared to nonmotor processing in VI/Crus I/Crus II and motor processing in I-VI. This hypothesis may be useful in the interpretation of future cerebellar neuroimaging findings. For example, this hypothesis may help interpret or highlight the potential relevance of isolated abnormalities in lobule VIII and IX in ADHD^97^. A virtue of this hypothesis is that it is testable using task fMRI. For example, future studies may contrast motor task conditions with high versus low task-focus demands (to isolate second motor representation), task-focused nonmotor task conditions with higher versus lower task-focus demands (to isolate third task-focused nonmotor representation), and task-unfocused nonmotor task conditions which can be removed from motor processing by, for example, modulating the amount of mental object manipulation to isolate the third task-unfocused nonmotor representation.

In sum, we describe a fundamental sensorimotor-fugal principle of organization in the cerebellum and highlight hierarchical similarities between cerebellar lobules VIII and IX/X. Our findings and analyses represent a significant conceptual advance in cerebellar systems neuroscience, and introduce novel approaches and testable questions to the investigation of cerebellar topography and function.

